## Supplementary Material for "Structural remodeling of target-SNARE protein complexes by NSF enables synaptic transmission"

Supplementary Figure 10 ........ 22

Supplementary Figure 11 ........ 28

Supplementary Table 1 ........ 29

Supplementary Table 2 .......................................................................................................................30

Supplementary Movie 1 .....................................................................................................................31

### Supplementary discussion

##### sx20S yield

Initially, a syntaxin concentration was selected for sx20S complex formation and cryo-EM based on the simplifying assumption of monomeric syntaxin in solution, and NSF:syntaxin:alpha-SNAP were mixed with the standard ratio of 1:5:30 as in the other experiments. Given the starting concentration of NSF (10 μM), the hypothetical concentration of monomeric syntaxin was 50 μM, somewhat close in hindsight to the reported oligomerization threshold determined previously by CD spectroscopy[1]. This led to a low yield of the sx20S complex, with around 5% of NSF forming sx20S complex with syntaxin and alpha-SNAP in the EM experiment. This demonstrates that tetrameric syntaxin was one of multiple syntaxin species in the original prep, and that the true concentration of tetramer was probably quite low; this is consistent with recent mass spectrometry analysis as well[2]. In summary, the low yield is sensible under the assumption that the soluble syntaxin preparation used was characterized by a concentration-dependent mixture of monomeric and oligomeric species, and that NSF and alpha-SNAP specifically engage parallel, tetrameric syntaxin oligomers.

Based on this reasoning, the syntaxin concentration was increased tenfold in subsequent assays for syntaxin tetramer disassembly and Munc18 capture to ensure concentrations exceeding the threshold for full oligomerization and to ensure that the absolute concentration of parallel tetramer was sufficiently high. This difference is evident upon comparison of Supplementary Figs. 2 and 3.

Conceptually, we note that *in vivo*, topography and the chemical environment of the membrane likely enrich for the formation of parallel, oligomeric species over the levels seen *in vitro* with soluble syntaxin. In particular, the transmembrane region of syntaxin ensures that it cannot fully sample rotational space, promoting parallel SNARE-SNARE interactions at the expense of antiparallel oligomeric syntaxin. Clustering related to interactions with membrane lipids or other factors would also enhance the effective local concentration of syntaxin and would act to drive it into the oligomeric state.

##### Comparison with p97

While this is the first structural evidence for the ADP•$P_i$ state in NSF, the mechanism is consistent with early biochemical study of myosin[3], which proposed the ADP•$P_i$ state and concerted release of $Mg^{2+}$ and $P_i$, as well as recent EM studies of the proteasome[4] and the NSF homolog p97[5] under hydrolyzing conditions. Indeed, during the preparation of this manuscript, a similar strategy was employed in the study of p97, albeit in the absence of a protein substrate[5]. Three key results from ref. [5] are of interest for comparison with our results.

First, Shein et al. note the presence of two distinct sites for $Mg^{2+}$ and $P_i$ following hydrolysis, termed states A and B. In state A, the $P_i$ engages in a bidentate interaction with the Sensor I residue N348 (N374 in NSF) and lies somewhat coaxial with the ADP phosphates, while $Mg^{2+}$

remains coordinated by the ADP Pβ and p97 Walker A residue T252 (T267 in NSF). In state B, both Pi and $Mg^{2+}$ shift downward, apparently related to rotameric shifts of the first arginine finger R359 (R385 in NSF) and a nearby phenylalanine, F360 (P386 in NSF). Furthermore, the p97 mutation F360P, which would remove this rotameric switching, is found to be hyperactivating. Strikingly, we only observe $Mg^{2+}$ and $P_i$ discretely bound in state A, and a proline residue is present at the position equivalent to F360 in wild-type NSF. This provides additional support to the suggestions made by Shein et al. that state B is a second transient state that follows state A after ~1 μs as assessed by molecular dynamics simulations, and opens the intriguing idea that the ATPase rate may be tuned by evolution through mutation at position 360/386 in p97/NSF. One might thus expect NSF P386F to have diminished activity and a detectable state B by cryo-EM.

Second, Shein et al. note large RMSF values for p97 MD simulations in the ATP state vs. the ADP•$P_i$ state, particularly at the Walker A helix and the NTD-D1 linker. While NSF does not show dramatic flipping of its N-domains with the transition from ATP to ADP, the N-D1 linker and nearby secondary structure elements are nevertheless differentially shifted in a nucleotide-dependent manner that appears propagated through the helix and down to the active site. Completely flipped N-domains are only observed in SNARE-substrate free NSF reconstructions (Supplementary Fig. 7).

Third, Shein et al. propose an inter-subunit communication pathway in p97. In both p97 and NSF, exactly 11 residues form a semi-structured link between the active sites of two protomers (the "sensor loop"), with the Sensor I motif at the N-terminal end and the first arginine finger at the C-terminal end. The sensor loop represents a plausible pathway by which the Sensor I motif of a given protomer can be modulated allosterically by the presence of nucleotide in the up-ring protomer, but evidence of any conformational transition here is absent in our datasets.

Finally, we do not observe evidence of large-scale conformational change within the loop of NSF corresponding to the so-called intersubunit signaling loop (ISS, NSF residues 359–364) in other AAA+ proteins, such as Yme1[6,7] and p97[8]. In both proteins, the ISS undergoes a dramatic, nucleotide-dependent transition, in which it is engaged in *trans* in the presence of up-ring ATP, and in *cis* in the presence of up-ring ADP. We observe local changes to the interprotomer interface accompanying $P_i$ release in NSF. However, these changes are limited to small-scale rotameric shifts; no dramatic conformational change is observed, even in the substrate-free, ADP-bound protomers. Given the high degree of sequence divergence in this region, the structural mechanism seen in Yme1 and p97 is likely not employed by NSF.

### Supplementary figures

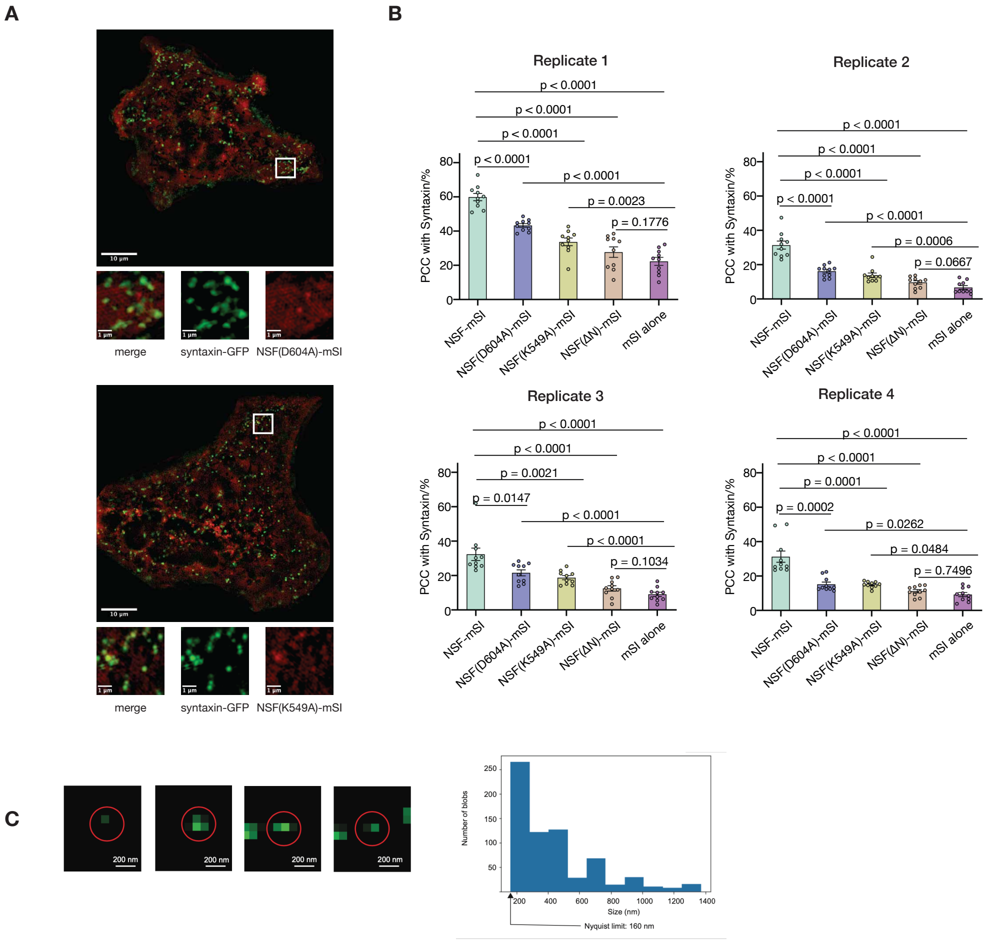

**Supplementary Figure 1 | Replicates and supporting materials for syntaxin colocalization**

**A.** Colocalization between syntaxin-GFP nanodomains and two NSF mutants, NSF(D604A)-mScarlet-I (PCC = 43% ± 1%, mean ± SEM) and NSF(K549A)-mScarlet-I (PCC = 33% ± 2%, mean ± SEM). NSF(D604A) is hydrolysis defective, while NSF(K549A) cannot bind nucleotide and thus fails to form a functional hexamer. **B.** Colocalization data and significance tests for four replicate imaging sessions of PC12 cells transfected with each of five different constructs. **C.** Size analysis of 20 SIM images of budding *S. cerevisiae* reveals discrete cluster sizes approaching the Nyquist limit; clusters are likely smaller.

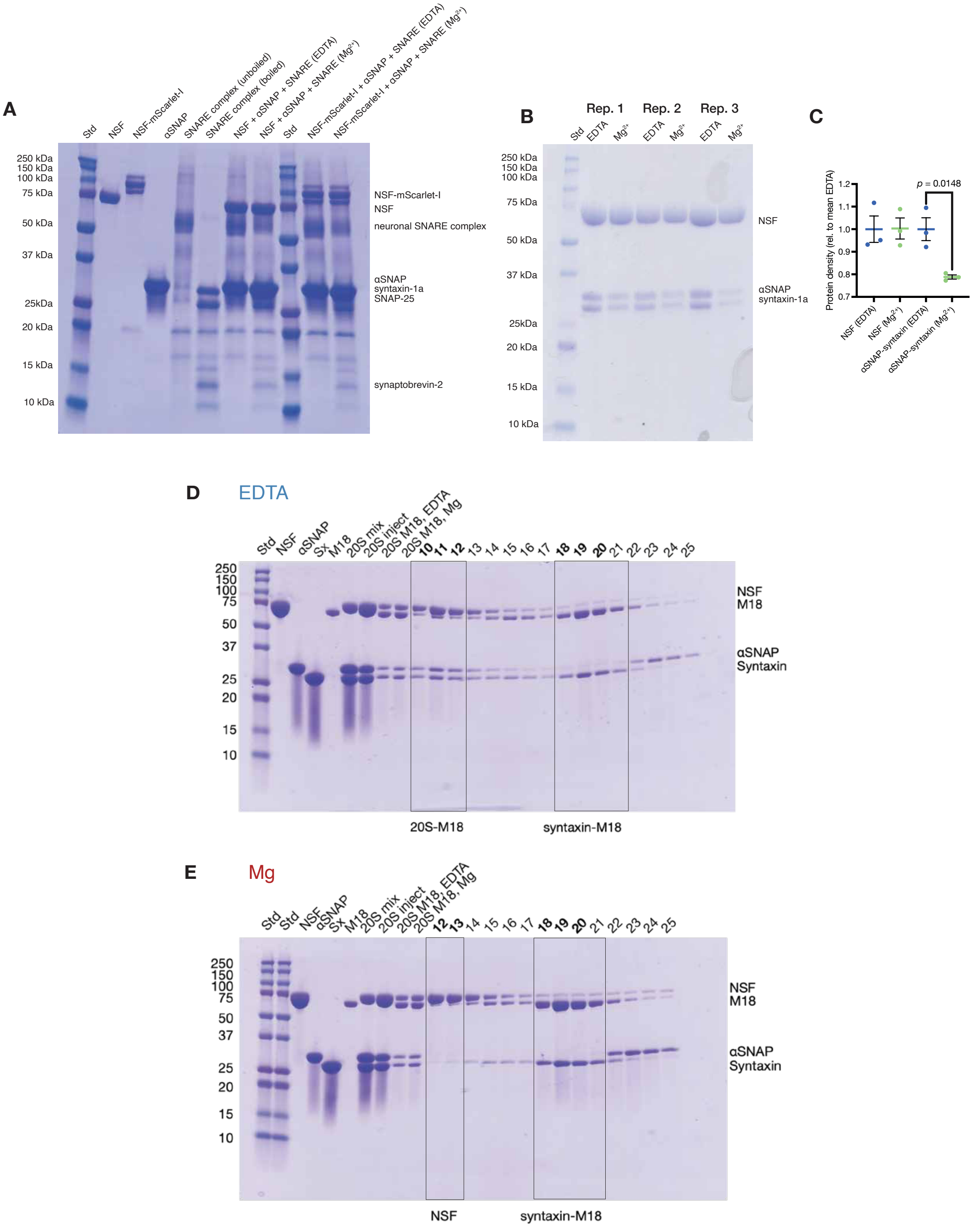

**Supplementary Figure 2 | Replicates and supporting materials for SNARE complex disassembly**

**A.** NSF-mScarlet-I retains its ability to disassemble the neuronal SNARE complex as determined by an electrophoretic mobility shift assay (EMSA). Individual components (NSF, NSF-mScarlet-I, α-SNAP, the neuronal SNARE complex) were purified independently (lanes 2–6) and mixed. Then, either EDTA or $Mg^{2+}$ were added as a control, or to initiate the disassembly of the SNARE complex, respectively (NSF, lanes 7 and 8; NSF-mScarlet-I, lanes 10 and 11). Note that the neuronal SNARE complex is SDS and heat resistant, and it runs as a single band (lane 5) unless boiled (lane 6) before SDS-PAGE; lanes 5 and 6 are thus fully assembled or disassembled controls. Comparison of EDTA and $Mg^{2+}$ lanes for either form of NSF shows the appearance of lower molecular weight species matching the boiled SNARE complex control, confirming the disassembly of the SNARE complex by each. NSF-mScarlet-I is thus expected to be hexameric and disassembly-competent in our cellular imaging experiments. **B, C.** NSF and α-SNAP disassemble oligomeric syntaxin as assessed by a gel-based de-enrichment assay. sx20S was purified and split into two pools. EDTA was added to the first pool as a non-hydrolyzing control, while $Mg^{2+}$ was added to the second to initiate syntaxin complex disassembly. Each pool was then concentrated using a 100,000 MWCO filtration device, retaining 20S complex or α-SNAP—syntaxin subcomplex but allowing individual syntaxin and α-SNAP molecules to pass, de-enriching them. Under hydrolyzing conditions, α-SNAP and syntaxin were significantly de-enriched by around 21 ± 1% ($p = 0.0148$) as assessed by SDS-PAGE, consistent with the disassembly of oligomeric syntaxin by NSF. **D, E.** SDS-PAGE corresponding to the SEC results shown in Fig. 1E. Samples corresponding to individual purified components, the 20S mixture pre-purification (20S mix), the purified 20S complex (20S inject), and the non-hydrolyzing (20S M18, EDTA) and hydrolyzing (20S M18, Mg) mixtures pre-injection as well as eluted fractions were run in both cases.

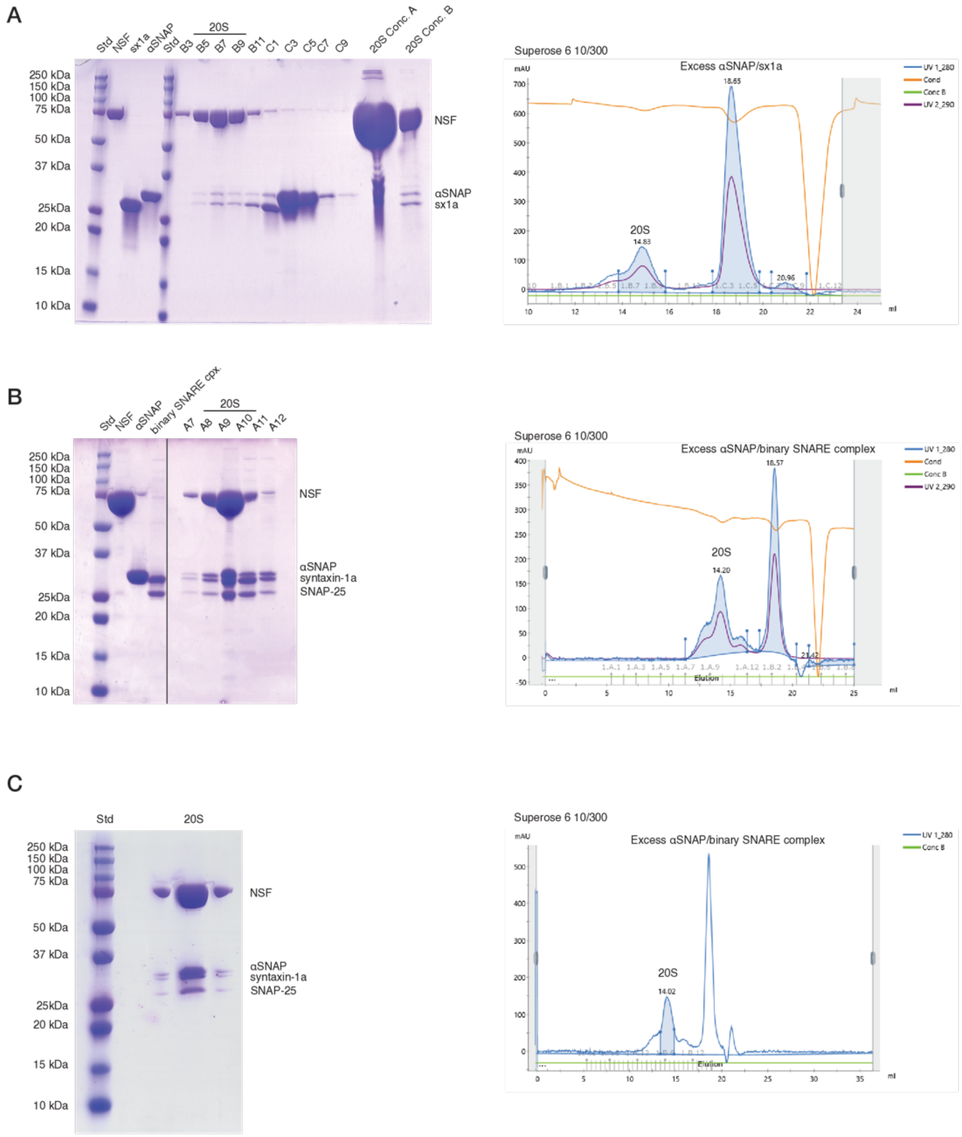

**Supplementary Figure 3 | Purification of various complexes for cryo-EM SPA**

NSF, the specified SNARE complexes, and α-SNAP were each purified recombinantly from *E. coli*, mixed with a 1:5:30 ratio (syntaxin tetramer) or a 1:3:15 ratio (binary complex) under non-hydrolyzing conditions, respectively, and subjected to SEC using a Superose 6 10/300 column. The initial concentration of NSF in each case was 10 µM and the injection volume was 500 µL. The specified complexes typically eluted with a retention volume of ~14.5 mL, were pooled generously, and were concentrated to ~15 mg/mL before freezing for cryo-EM. SDS-PAGE was subsequently used to verify the presence of all proteins in the complex. **A.** SDS-PAGE and SEC of the sx20S complex comprised of NSF, α-SNAP, and soluble syntaxin. EDTA was included throughout to chelate any divalent cation. **B.** SDS-PAGE and SEC of the 21bin20S complex comprised of NSF, α-SNAP, and the syntaxin—SNAP-25 complex. It comprises two soluble syntaxin molecules and one full-length SNAP-25 (2:1 complex). EDTA was included throughout to chelate any divalent cation. **C.** SDS-PAGE and SEC of the 21bin20S complex comprised of NSF, α-SNAP, and the syntaxin—SNAP-25 complex. This stage consists of two soluble syntaxin molecules and one full-length SNAP-25. As in the case of the preparation under non-hydrolyzing conditions, divalent cation was not added to the SEC buffer. However, EDTA was not included, and purification was performed quickly. Following gel filtration and concentration, $MgCl_2$ was added to start the disassembly reaction, converting the sample to the 2:2 complex.

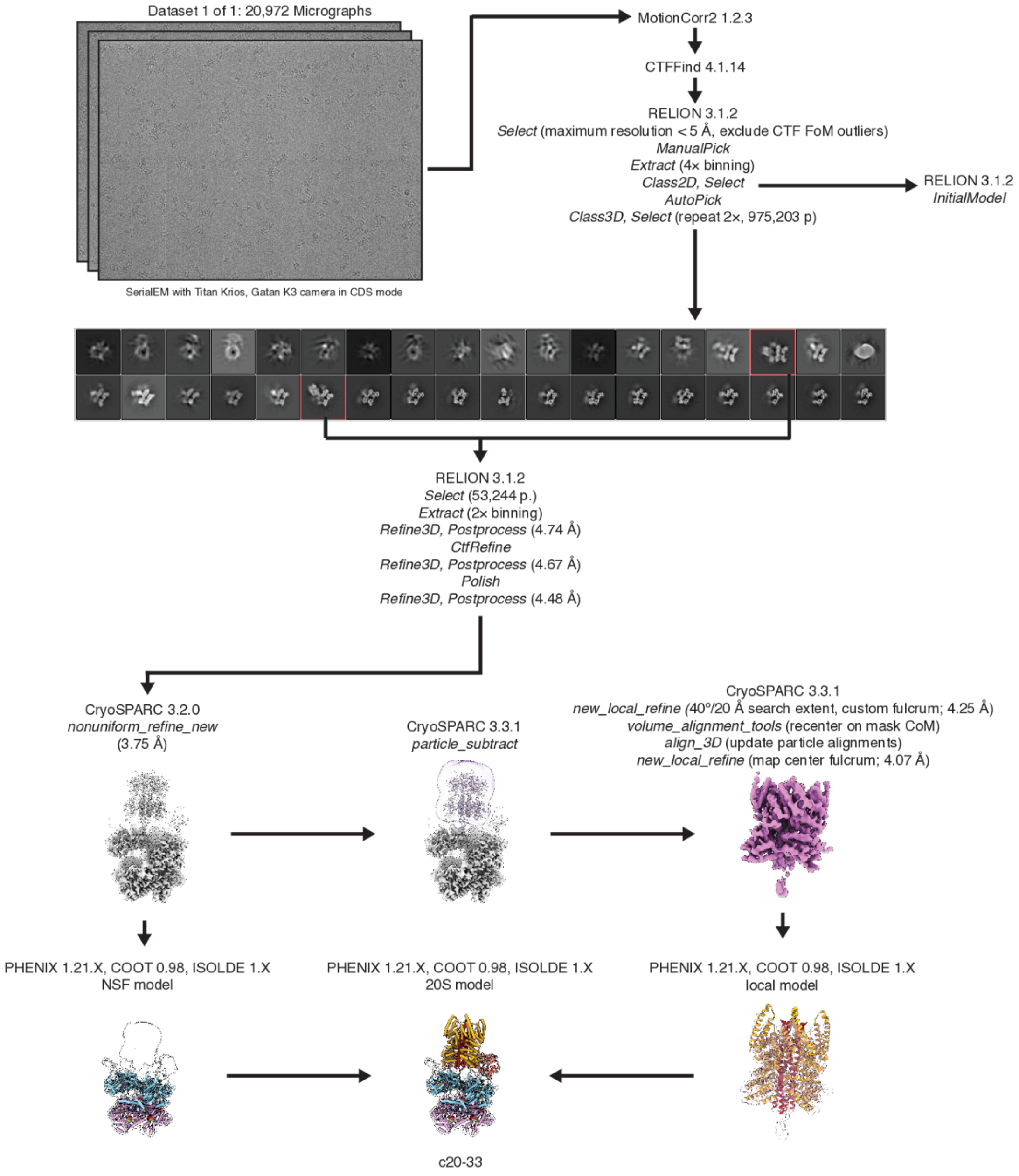

**Supplementary Figure 4 | Cryo-EM SPA data collection and processing workflow for the sx20S complex**

A single sx20S cryo-EM dataset frozen under non-hydrolyzing conditions and comprised of 20,972 micrographs was collected and processed, yielding a single sx20S class with 53,244 particles. The remaining classes were substrate-free and were discarded; this is not due to the disassembly of the syntaxin tetramer, but rather the inability to fully separate NSF from the sx20S complex by SEC. The sx20S particles produced a single sx20S reconstruction with an overall resolution of 3.75 Å. Modeling of the NSF D1 and D2 rings was performed with this reconstruction. Subsequently, this reconstruction was used as the basis for signal subtraction and local reconstruction of the α-SNAP—syntaxin tetramer subcomplex, and the α-SNAP—syntaxin tetramer subcomplex model was built with this reconstruction and then docked back into the full sx20S reconstruction. See Methods for further details.

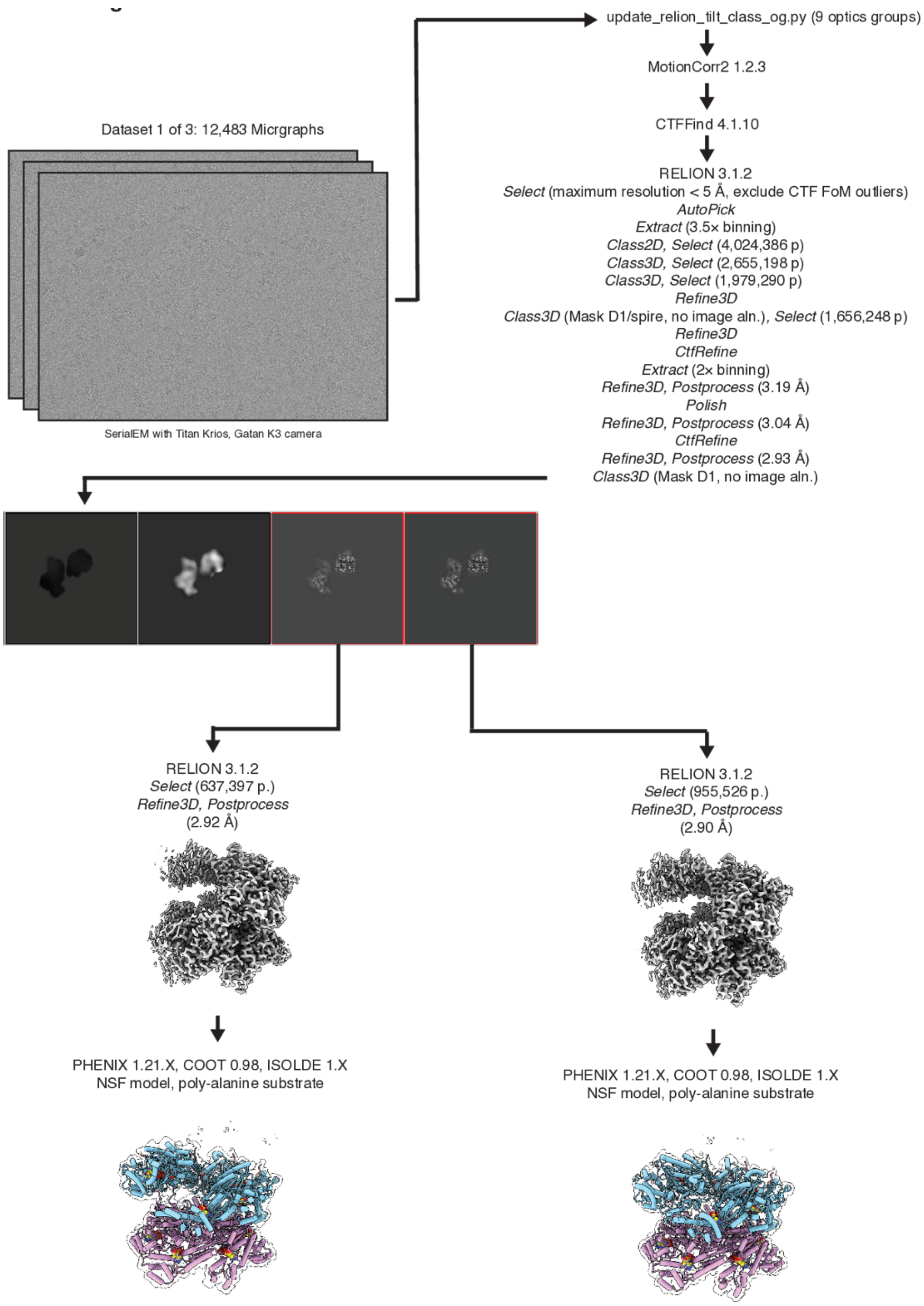

**Supplementary Figure 5 | Cryo-EM SPA data collection and processing workflow for the high-resolution NSF D1/D2 reconstructions under non-hydrolyzing conditions**

A single cryo-EM SPA dataset from 21bin20S sample frozen under non-hydrolyzing conditions and comprised of 12,483 micrographs was collected and processed, yielding two classes (class 4-3-5-3, engaged F protomer, 2.92 Å, and class 4-3-5-4, disengaged F protomer, 2.90 Å) with near-atomic resolution D1 and D2 reconstructions. The data processing strategy averages out spire density and thus precludes α-SNAP—SNARE subcomplex modeling, so substrate in the D1 pores of each class is modeled as a poly-alanine chain. Due to their high quality, these models served as starting points for all other classes reported herein. See Methods for further details.

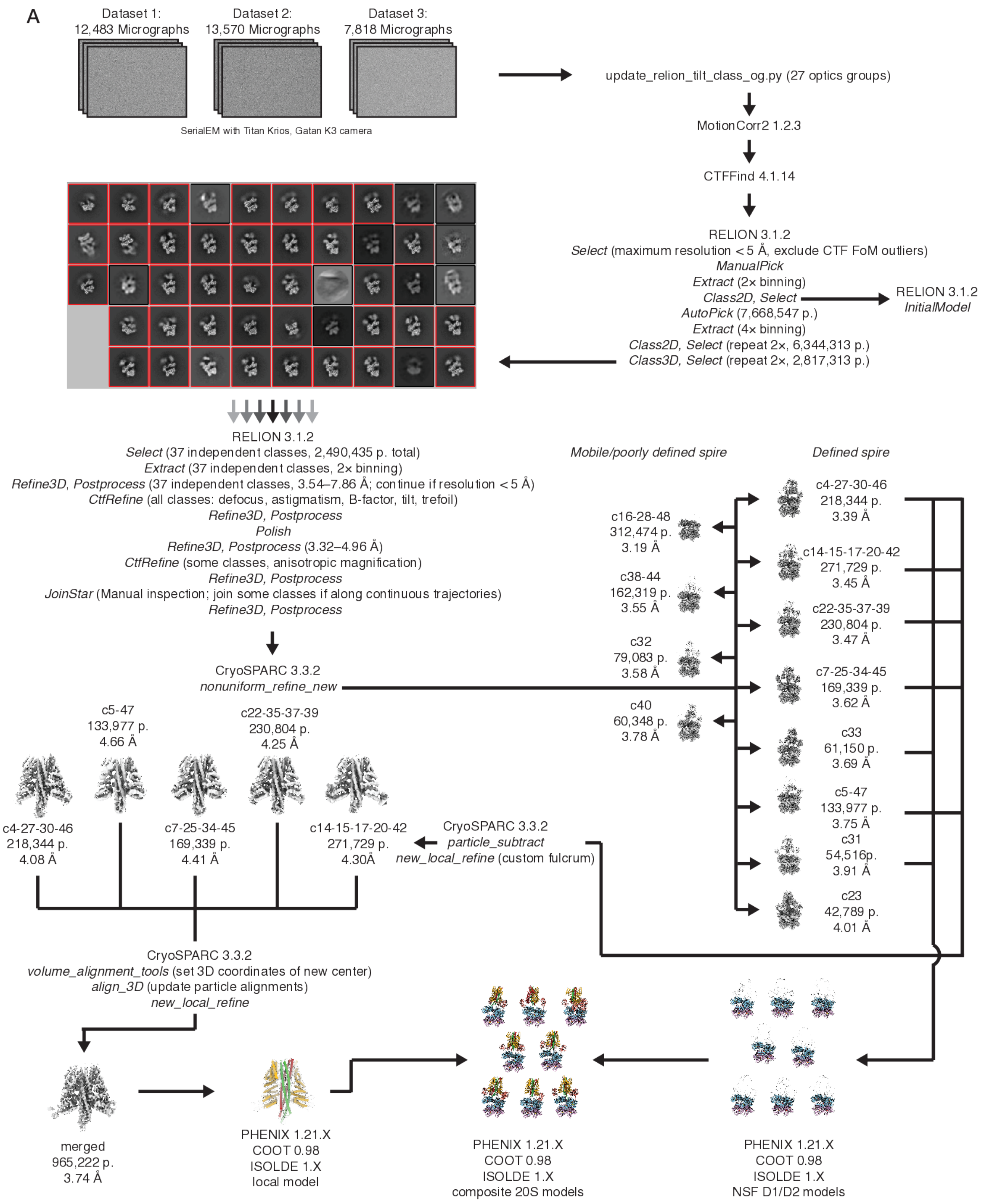

**Supplementary Figure 6 | Cryo-EM SPA data collection and processing workflow for the 21bin20S complex under non-hydrolyzing conditions**

Three cryo-EM SPA datasets collected from the 21bin20S sample frozen under non-hydrolyzing conditions, including the dataset described in Supplementary Fig. 5, were combined for a total of 33,871 micrographs. Ultimately, nearly 2.5 million particles were classified into 37 independent classes; classes along continuous trajectories of conformational change were merged, yielding twelve total classes with resolutions ranging from 3.19–4.01 Å. Eight classes with at least partially ordered spire density were carried forward to model the NSF D1 and D2 ATPase rings; all show substrate-engaged F protomers and differ largely based on spire pose. Signal subtraction and local refinement of the α-SNAP—SNARE subcomplex of five of these classes having more than 100,000 particles was then performed. Then, the particles from these local reconstructions were subsequently aligned to yield one final merged reconstruction with 965,222 particles and a final resolution of 3.74 Å. A model was built into the α-SNAP—2:1 binary SNARE subcomplex and subsequently docked back into the full 21bin20S reconstructions, yielding eight total 21bin20S models. See Methods for more details.

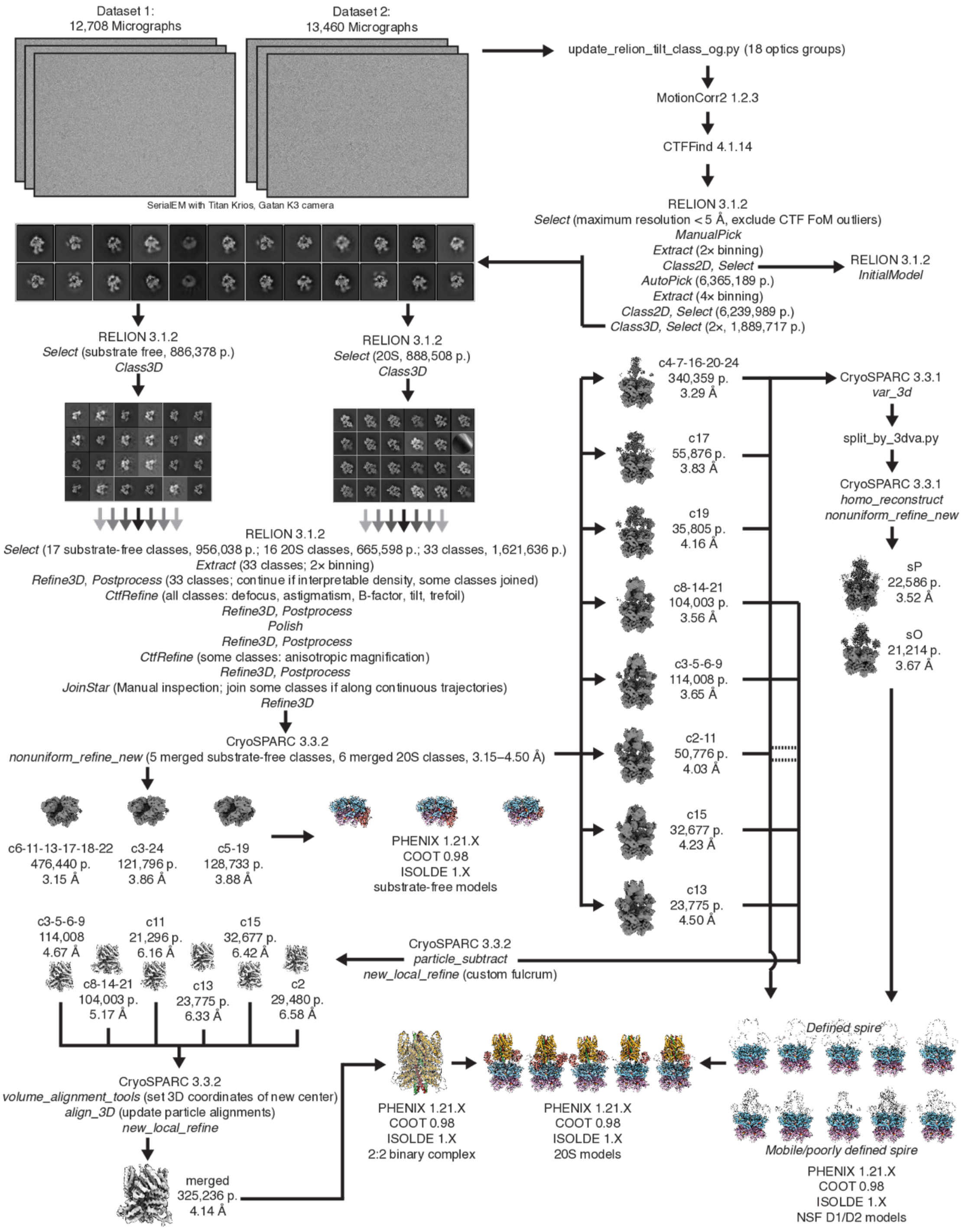

**Supplementary Figure 7 | Cryo-EM SPA data collection and processing workflow for the 22bin20S complex under hydrolyzing conditions**

Two cryo-EM SPA datasets were collected from 22bin20S samples frozen under hydrolyzing conditions ~1 min after adding a mixture containing $Mg^{2+}$ and ATP regeneration components creatine phosphate and creatine kinase. Together, these data collections yielded 26,168 total micrographs. 3D classification of nearly two million particles yielded classes falling into two categories—22bin20S, with ordered α-SNAP—SNARE subcomplex density and density in the NSF D1 ATPase pore, or substrate-free NSF, with no density in the pore and a distinctive double split in both the D1 and D2 rings. After splitting these ~1.6 million particles into roughly equal sets, performing further independent 3D classifications, and merging classes with continuous disorder, a total of eight 22bin20S classes with ~600,000 particles (3.29–4.50 Å) and three substrate-free NSF classes with ~1 million particles (3.15–3.88 Å) were carried forward. 3DVA was performed on the largest 22bin20S class (class 4-7-16-20-24), which revealed substantial conformational change in the NSF D1 ring over the first mode. Around 20,000 particles from the tails of the 3DVA distribution were thus used to produce two additional 22bin20S reconstructions, for a total of ten. Models were then built into the three substrate-free NSF reconstructions and the ten 22bin20S reconstructions. Signal subtraction and local refinement of the α-SNAP—SNARE subcomplex of six 22bin20S classes were then performed. Then, the particles from these local reconstructions were subsequently aligned to yield one final merged reconstruction with 325,236 particles and a final resolution of 4.14 Å. A model was built into the α-SNAP—2:2 binary SNARE subcomplex and subsequently docked back into the full 22bin20S reconstructions with well-defined spires, yielding five total 22bin20S models. See Methods for more details.

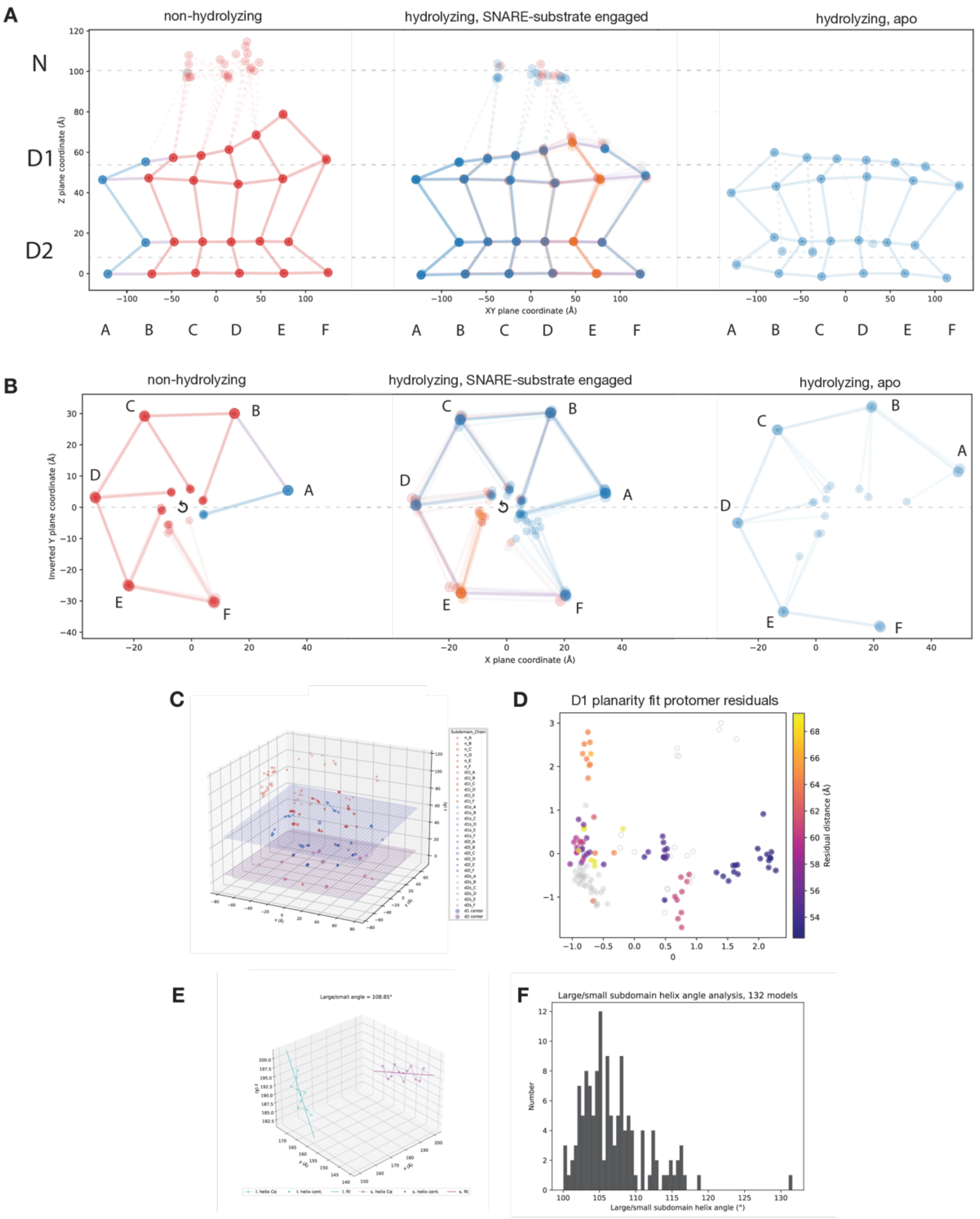

**Supplementary Figure 8 | Additional details for NSF D1 and D2 planarity analysis and large-small subdomain angle analysis.**

These panels supplement Fig. 5. **A.** Centers of mass for N-domains and the D1 large, D1 small, D2 large, and D2 small subdomains in 3D and planes were fit to the subdomains corresponding to the D1 and D2 rings (see also **C**). The results are projected into the 2D, with the NSF protomer subdomain centers of mass unrolled into a single plane to illustrate their relative positions. Dashed lines indicate the plane fit. A given protomer is colored by the nucleotide found in its D1 subdomain; ADP-bound protomers are colored blue, ATP-bound protomers are colored red, ATP-$Mg^{2+}$-bound protomers are colored orange, and nucleotide-free protomers are colored grey. This comparison illustrates the high degree of structural similarity between the D2 rings between datasets, in contrast to the high degree of conformational change in the up-ring protomers of D1. These conformational differences are largely coaxial with the D1 pore. More variation is present in the hydrolyzing dataset than in the non-hydrolyzing dataset. **B.** Projection of D1 protomer centers of mass into the plane of the D1 ring reveals conformational change associated with protomer F rebinding; the non-hydrolyzing structure projection reveals the limits of F protomer disengagement and engagement, while protomers F in the hydrolyzing dataset are more variable and transition between these states. This projection also illustrates how the SNARE substrate-free D1 rings retract from the pore axis. **C.** Raw plane fits and centroids in 3D used to generate **A** and **B**. See Methods for more details. **D.** NSF large subdomain and small subdomain D1 plane fit residuals for a given protomer (substrate-bound, hydrolyzing dataset protomers A–F and non-hydrolyzing protomers A and F) were summed and used to color structures projected into the PCA space (Fig. 8) to illustrate planarity changes along the conformational coordinate associated with hydrolysis. **E.** Example helical axes fits for one of 132 NSF D1 large-small subdomain angle calculations. The smallest angle between vectors defined by these fits is then calculated. **F**. Overall distribution of large-small subdomain angles over 132 protomers from 22 models in the dataset.

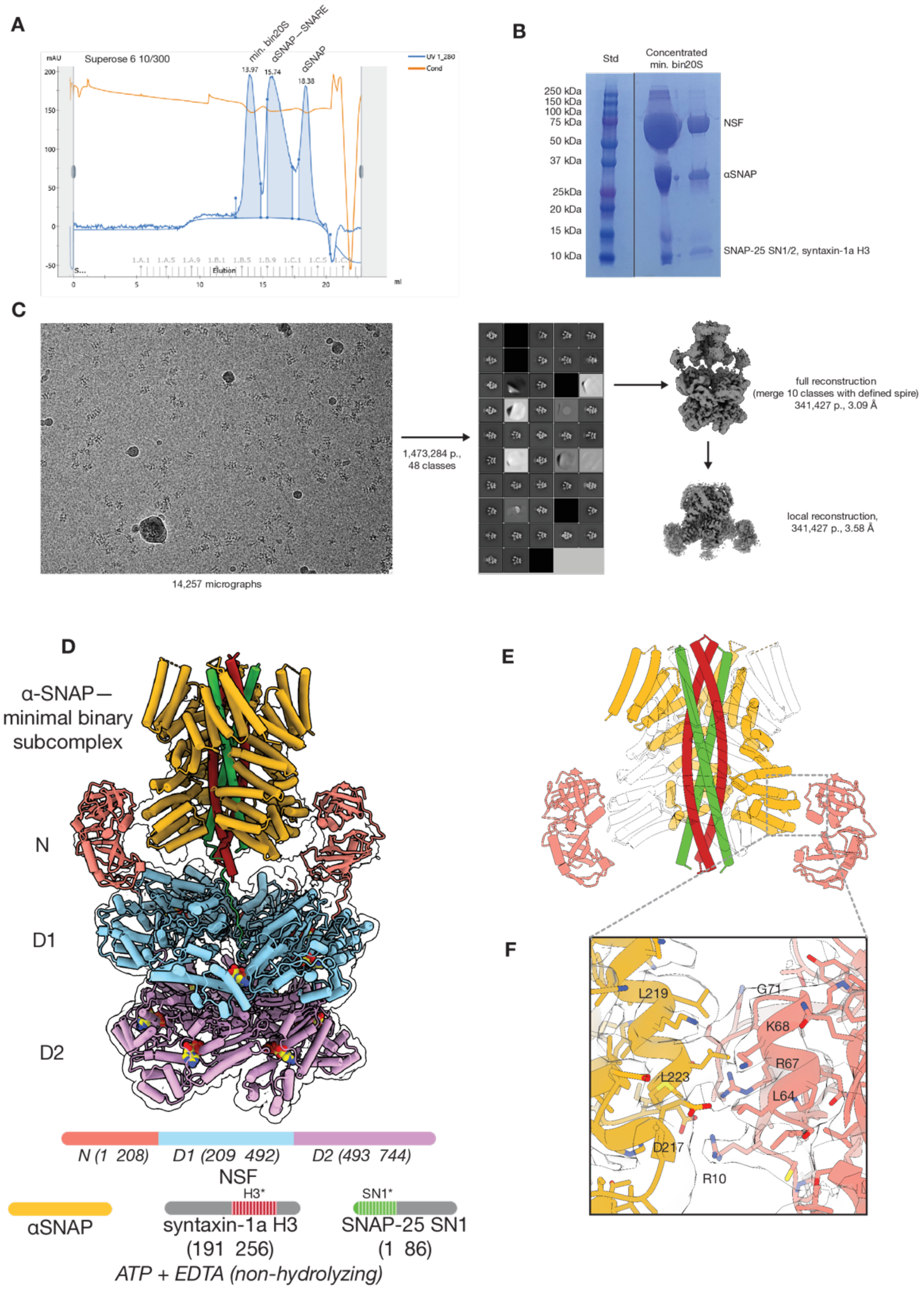

**Supplementary Figure 9 | NSF—α-SNAP—2:2 minimal syntaxin—SNAP-25 complex structure**

**A.** NSF, α-SNAP, SNAP-25A SN1 SNARE domain (residues -2–83), SNAP-25A SN2 SNARE domain (residues 141–204), and syntaxin-1a H3 SNARE domain were each purified recombinantly from *E. coli*, mixed with a 1:5:20 ratio under non-hydrolyzing conditions, respectively, and subjected to size exclusion chromatography (SEC) using a Superose 6 10/300 column. The minimal bin20S complex eluted with a retention volume of ~14 mL and was concentrated to ~20 mg/mL before freezing for cryo-EM. SDS-PAGE was subsequently used to verify the presence of the 20S complex. **B.** A single minimal bin20S cryo-EM dataset frozen under non-hydrolyzing conditions and comprised of 14,257 micrographs was collected and processed. As the primary goal of this collection was to identify the stoichiometry of the SNARE substrate in the spire, all classes with defined spire density were merged into a single minimal bin20S superclass with 341,427 particles. These particles were used to reconstruct density for the minimal bin20S complex with a resolution of 3.09 Å. Modeling of the NSF D1 and D2 rings was performed with this reconstruction. Subsequently, this reconstruction was used as the basis for signal subtraction and local reconstruction of the NSF N-domain—α-SNAP—binary SNARE subcomplex to 3.86 Å resolution. **C.** The overall architecture of the minimal bin20S supercomplex. **D.** The structure of the minimal 2:2 syntaxin—SNAP-25 subcomplex (class #27), with two of four α-SNAP molecules hidden for clarity. This configuration matches a previously reported crystal structure of the binary complex[10] and the full-length bin20S SNARE subcomplex observed by cryo-EM in this work (Fig. 4). While the SNAP-25 SN2 SNARE domain was copurified with the other two SNARE domains, it was not present in the EM density. SNARE domains included in the model are indicated with dashed lines and an asterisk. **E.** The local reconstruction of the α-SNAP—minimal 2:2 syntaxin—SNAP-25 subcomplex (class #28) reveals new detail into the primary interaction between the N-domains and α-SNAP. This interface is characterized by hydrophobic α-SNAP sidechains L219 and L223 packing near NSF N-domain residues 64–71; both sides of this interaction are α-helical. Nearby, α-SNAP D217 forms a salt bridge with NSF N-domain R10.

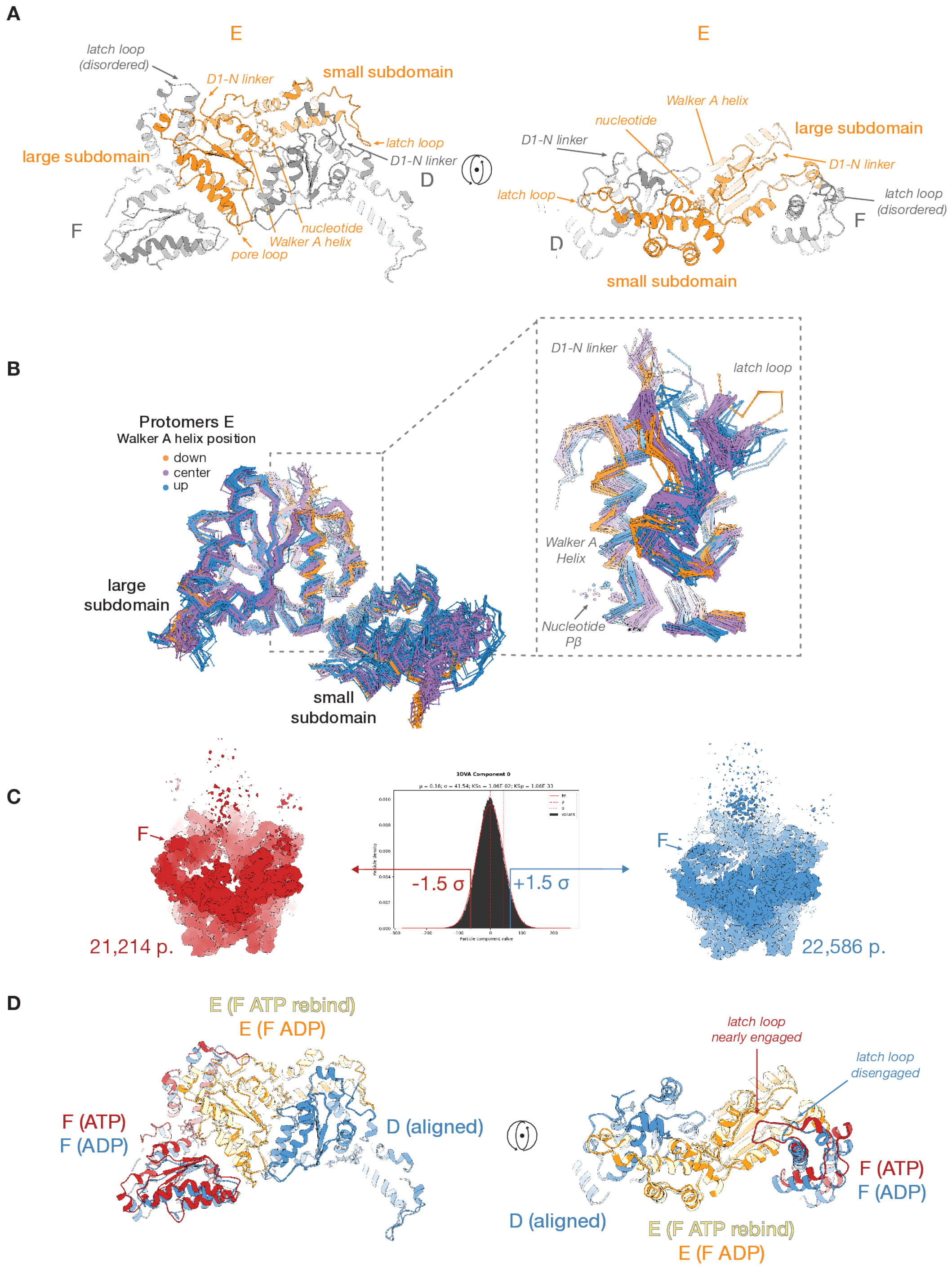

**Supplementary Figure 10 | Unique features of protomers E from the hydrolyzing dataset are connected to protomer F rebinding.**

**A.** Two views of a protomer E from the hydrolyzing dataset (colored orange), with neighboring protomers F and D (colored grey). Various notable features are annotated. **B.** All protomers from the hydrolyzing condition, and protomers A and F from the non-hydrolyzing condition are aligned by their D1 large subdomains. SNARE substrate-free models from the hydrolyzing condition are not shown. N-domains and D2 domains are hidden. Protomers E from the hydrolyzing condition show a unique Walker A α-helix conformation (orange models, “down”), in which the helix is shifted down and tilted towards the D1 pore. Most protomers fall into an intermediate state (purple, “center”), except for the protomers A, which occupy a third state (blue, “up”). This is connected to two key features—the N-D1 linker position, for which protomers E have a unique conformation, and the latch loop conformation, which varies from a disengaged and helical state, through an unstructured state, and ultimately to an engaged state. In all but one case, protomers E have a disengaged protomer F up-ring, and thus an intermediate latch loop state. Critically, the engagement of protomer F involves an interaction between the latch loop and the N-D1 linker, which is pulled away from the Walker A helix, allowing it to shift into the center or up states (inset, right). **C.** *CryoSPARC* 3D Variability Analysis (3DVA)[9] captured the transition between the disengaged and engaged protomer F states. 3DVA was run on a large class from the hydrolyzing dataset, and particles in the ±1.5 σ tails of the first mode were used to reconstruct the most extreme states along the coordinate. The change in protomer F state, as well as a shift in protomer A towards the split, are evident on inspection of unsharpened maps. **D.** Sub-models showing the states of protomers D, E, and F derived from the density in **C**. The latch loop of protomer F engages protomer E to a degree approaching the state found in the non-hydrolyzing condition (not shown).

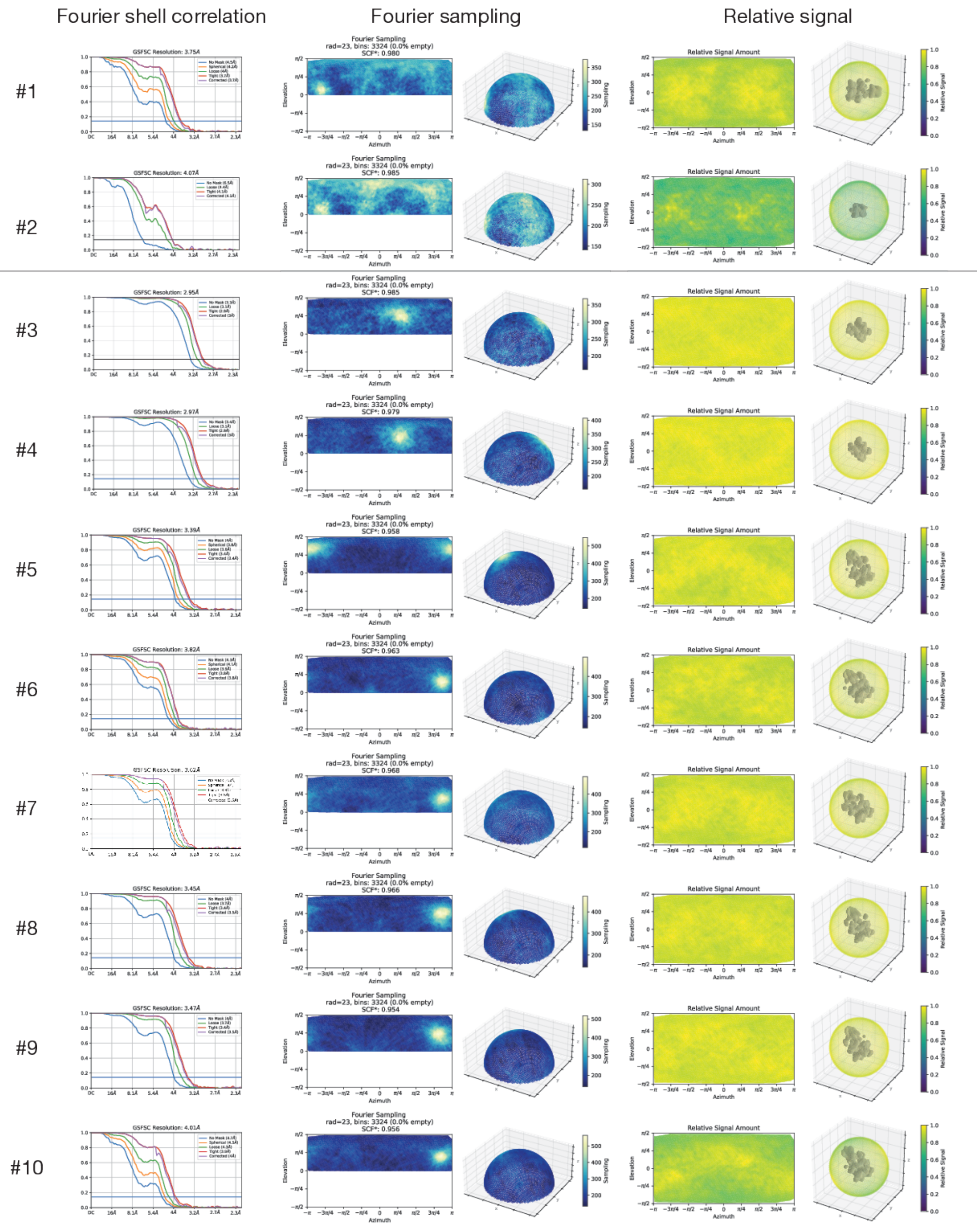
Fourier shell correlation
Fourier sampling
Relative signal
#1
#2
#3
#4
#5
#6
#7
#8
#9
#10

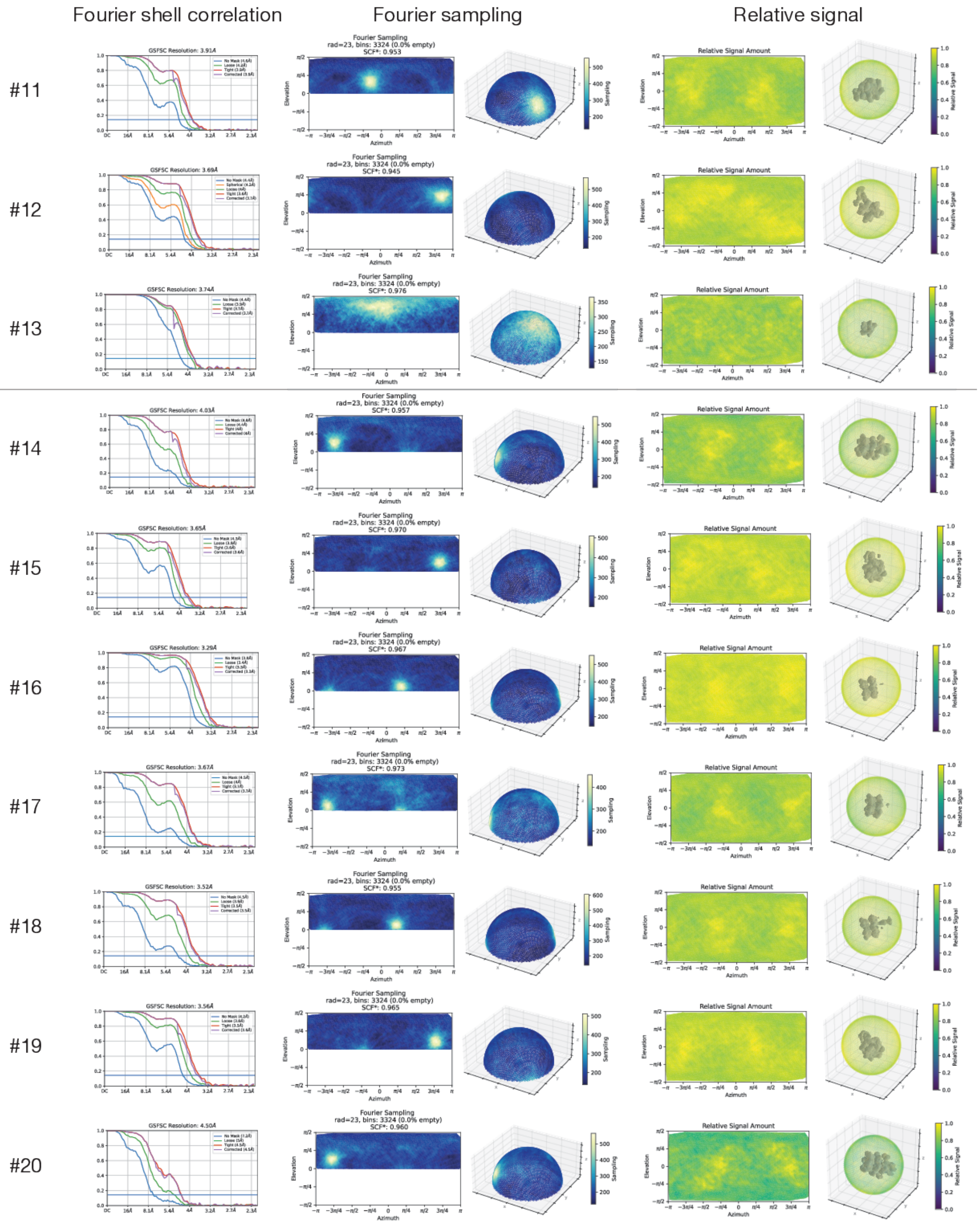

Fourier shell correlation
Fourier sampling
Relative signal
#11
#12
#13
#14
#15
#16
#17
#18
#19
#20
GSFSC Resolution: 3.91Å
GSFSC Resolution: 3.69Å
GSFSC Resolution: 3.74Å
GSFSC Resolution: 4.03Å
GSFSC Resolution: 3.65Å
GSFSC Resolution: 3.29Å
GSFSC Resolution: 3.67Å
GSFSC Resolution: 3.52Å
GSFSC Resolution: 3.56Å
GSFSC Resolution: 4.50Å
Fourier Sampling
rad=23, bins: 3324 (0.0% empty)
SCF*: 0.953
SCF*: 0.945
SCF*: 0.976
SCF*: 0.957
SCF*: 0.970
SCF*: 0.967
SCF*: 0.973
SCF*: 0.955
SCF*: 0.965
SCF*: 0.960
Relative Signal Amount
Elevation
Azimuth
Sampling
Relative Signal

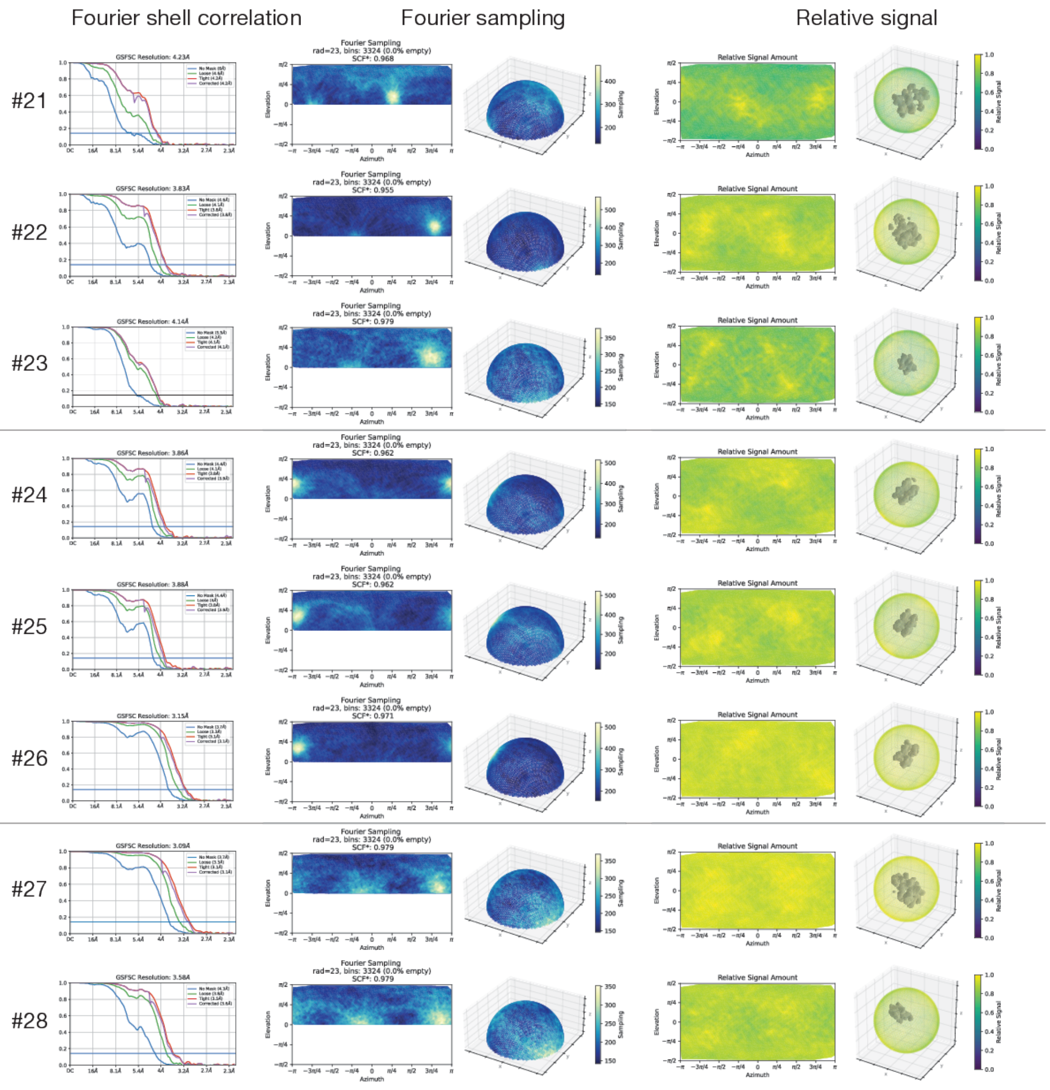

**Supplementary Figure 11 | Fourier shell correlation (FSC) curves, Fourier sampling, and relative signal for all reconstructions.**

See Tables S1–2 for corresponding details.

| | NSF—αSNAP—syntaxin H3 SNARE domain tetramer (substrate bound, non-hydrolyzing; sx20S) | | NSF—αSNAP—2:1 syntaxin:SNAP-25 SNARE complex (substrate bound, non-hydrolyzing; 21bin20S) | | | | | | | | | | |
|---|---|---|---|---|---|---|---|---|---|---|---|---|---|
| | #1 (c20-33 full) | #2 (c20-33 local) | #3 (c4-3-5-3) | #4 (c4-3-5-4) | #5 (c4-27-30-46) | #6 (c5-47) | #7 (c7-25-34-45) | #8 (c14-15-20-42 ) | #9 (c22-35-37-39 ) | #10 (c23) | #11 (c31) | #12 (c33) | #13 (merge, local) |
| | (EMDB-XXXXX) | (EMDB-XXXXX) | (EMDB-XXXXX) | (EMDB-XXXXX) | (EMDB-XXXXX) | (EMDB-XXXXX) | (EMDB-XXXXX) | (EMDB-XXXXX) | (EMDB-XXXXX) | (EMDB-XXXXX) | (EMDB-XXXXX) | (EMDB-XXXXX) | (EMDB-XXXXX) |
| | (PDB YYYY) | (PDB YYYY) | (PDB YYYY) | (PDB YYYY) | (PDB YYYY) | (PDB YYYY) | (PDB YYYY) | (PDB YYYY) | (PDB YYYY) | (PDB YYYY) | (PDB YYYY) | (PDB YYYY) | (PDB YYYY) |
| **Data collection and processing** | | | | | | | | | | | | | |
| Microscope, camera | Titan Krios, K3 | Titan Krios, K3 | Titan Krios, K3 | Titan Krios, K3 | Titan Krios, K3 | Titan Krios, K3 | Titan Krios, K3 | Titan Krios, K3 | Titan Krios, K3 | Titan Krios, K3 | Titan Krios, K3 | Titan Krios, K3 | Titan Krios, K3 |
| Magnification | 22,500× | 22,500× | 22,500× | 22,500× | 22,500× | 22,500× | 22,500× | 22,500× | 22,500× | 22,500× | 22,500× | 22,500× | 22,500× |
| Voltage (kV) | 300 | 300 | 300 | 300 | 300 | 300 | 300 | 300 | 300 | 300 | 300 | 300 | 300 |
| Dataset ID(s) | 20210903 | 20210903 | 20190624 | 20190624 | 20190624, 20190919, 20191216 | 20190624, 20190919, 20191216 | 20190624, 20190919, 20191216 | 20190624, 20190919, 20191216 | 20190624, 20190919, 20191216 | 20190624, 20190919, 20191216 | 20190624, 20190919, 20191216 | 20190624, 20190919, 20191216 | 20190624, 20190919, 20191216 |
| Electron exposure (e–/Å²) | 27.608 | 27.608 | 31.560 | 31.560 | 31.560, 36.760, 35.480 | 31.560, 36.760, 35.480 | 31.560, 36.760, 35.480 | 31.560, 36.760, 35.480 | 31.560, 36.760, 35.480 | 31.560, 36.760, 35.480 | 31.560, 36.760, 35.480 | 31.560, 36.760, 35.480 | 31.560, 36.760, 35.480 |
| Defocus range (μm) | -1.0 to -3.0 | -1.0 to -3.0 | -1.0 to -3.0 | -1.0 to -3.0 | -1.0 to -3.0 | -1.0 to -3.0 | -1.0 to -3.0 | -1.0 to -3.0 | -1.0 to -3.0 | -1.0 to -3.0 | -1.0 to -3.0 | -1.0 to -3.0 | -1.0 to -3.0 |
| Physical pixel size (Å) | 1.096 | 1.096 | 1.096 | 1.096 | 1.096 | 1.096 | 1.096 | 1.096 | 1.096 | 1.096 | 1.096 | 1.096 | 1.096 |
| Symmetry imposed | None (C1) | None (C1) | None (C1) | None (C1) | None (C1) | None (C1) | None (C1) | None (C1) | None (C1) | None (C1) | None (C1) | None (C1) | None (C1) |
| Initial particle images (no.) | 975203 | 975203 | 1656248 | 1656248 | 2817313 | 2817313 | 2817313 | 2817313 | 2817313 | 2817313 | 2817313 | 2817313 | 2817313 |
| Final particle images (no.) | 53244 | 53244 | 637397 | 955526 | 218344 | 133977 | 169339 | 271729 | 230804 | 42789 | 54516 | 61150 | 965222 |
| Map resolution (Å) FSC threshold = 0.143 | 3.75 | 4.07 | 2.95 | 2.97 | 3.39 | 3.82 | 3.62 | 3.45 | 3.47 | 4.01 | 3.91 | 3.69 | 3.74 |
| Map sharpening B factor (Å²) | 117.0 | 148.3 | -80.8 | -70.77 | 129.8 | 145.2 | 141.2 | 119.6 | 145.9 | 114.2 | 121.7 | 112.3 | 221.1 |
| **Model Refinement** | | | | | | | | | | | | | |
| Initial model used (PDB ID) | #2–4 | 1N7S | 6MDM | 6MDM | #3–4, #13 | #3–4, #13 | #3–4, #13 | #3–4, #13 | #3–4, #13 | #3–4, #13 | #3–4, #13 | #3–4, #13 | 1N7S |
| Model composition | | | | | | | | | | | | | |
| *Non-hydrogen atoms* | 40730 | 10949 | 26375 | 26521 | 42082 | 37311 | 38879 | 41286 | 40952 | 43495 | 37457 | 39769 | 8953 |
| *Protein residues* | 5133 | 1377 | 3195 | 3190 | 5291 | 4690 | 4906 | 5100 | 5068 | 5466 | 4673 | 5027 | 1130 |
| *Ligands* | 13 | 0 | 12 | 11 | 12 | 11 | 12 | 12 | 12 | 11 | 11 | 12 | 0 |
| B factors (Å²) | | | | | | | | | | | | | |
| *Protein* | 122.90 | 105.57 | 46.17 | 180.56 | 287.92 | 82.07 | 159.80 | 76.07 | 69.85 | 153.83 | 93.84 | 50.83 | 96.20 |
| R.M.S. deviations | | | | | | | | | | | | | |
| *Bond lengths (Å)* | 0.007 | 0.0026 | 0.004 | 0.004 | 0.005 | 0.005 | 0.005 | 0.004 | 0.004 | 0.004 | 0.004 | 0.007 | 0.004 |
| *Bond angles (°)* | 1.220 | 0.55 | 0.699 | 0.799 | 1.002 | 1.018 | 1.066 | 0.947 | 0.974 | 0.976 | 0.996 | 1.116 | 0.693 |
| Validation | | | | | | | | | | | | | |
| *MolProbity score* | 1.40 | 1.34 | 1.04 | 1.31 | 1.40 | 1.32 | 1.31 | 1.38 | 1.43 | 1.54 | 1.69 | 1.41 | 1.50 |
| *Clashscore* | 3.58 | 4.67 | 2.37 | 4.69 | 4.82 | 4.80 | 3.61 | 3.97 | 5.12 | 6.78 | 8.01 | 4.94 | 8.23 |
| *Rotamer outliers (%)* | 1.03 | 0 | 0.76 | 0.75 | 0.22 | 0.75 | 1.08 | 0.25 | 0.30 | 0.62 | 1.34 | 0.58 | 0.74 |
| Ramachandran Statistics | | | | | | | | | | | | | |
| *Favored (%)* | 96.31 | 97.50 | 97.92 | 97.67 | 97.20 | 97.68 | 97.29 | 96.76 | 97.11 | 97.05 | 97.09 | 97.19 | 97.75 |
| *Allowed (%)* | 3.65 | 2.42 | 1.98 | 2.21 | 2.76 | 2.28 | 2.67 | 3.14 | 2.77 | 2.82 | 2.59 | 2.79 | 2.16 |
| *Outliers (%)* | 0.04 | 0.07 | 0.09 | 0.13 | 0.04 | 0.04 | 0.04 | 0.10 | 0.12 | 0.13 | 0.32 | 0.02 | 0.09 |

| | NSF—αSNAP—2:2 syntaxin:SNAP-25 SNARE complex (substrate bound, hydrolyzing; 22bin20S) | | | | | | | | | | NSF (substrate free, hydrolyzing) | | | NSF—αSNAP—minimal 2:2 syntaxin:SNAP-25 SNARE complex (substrate bound, hydrolyzing; min22bin20S) | |
|---|---|---|---|---|---|---|---|---|---|---|---|---|---|---|---|
| | #14 (c2-11) | #15 (c3-5-6-9) | #16 (c4-7-16-20-2 4) | #17 (c4-7-16-20-2 4-m0-sO) | #18 (c4-7-16-20-2 4-m0-sP) | #19 (c8-14-21) | #20 (c13) | #21 (c15) | #22 (c17) | #23 (merge, local) | #24 (c3-24) | #25 (c5-19) | #26 (c6-XXX-22) | #27 (c5-XXX-46) | #28 (c5-XXX-46) |
| | (EMDB-XXXXX) | (EMDB-XXXXX) | (EMDB-XXXXX) | (EMDB-XXXXX) | (EMDB-XXXXX) | (EMDB-XXXXX) | (EMDB-XXXXX) | (EMDB-XXXXX) | (EMDB-XXXXX) | (EMDB-XXXXX) | (EMDB-XXXXX) | (EMDB-XXXXX) | (EMDB-XXXXX) | (EMDB-XXXXX) | (EMDB-XXXXX) |
| | (PDB YYYY) | (PDB YYYY) | (PDB YYYY) | (PDB YYYY) | (PDB YYYY) | (PDB YYYY) | (PDB YYYY) | (PDB YYYY) | (PDB YYYY) | (PDB YYYY) | (PDB YYYY) | (PDB YYYY) | (PDB YYYY) | (PDB YYYY) | (PDB YYYY) |
| **Data collection and processing** | | | | | | | | | | | | | | | |
| Microscope, camera | Titan Krios, K3 | Titan Krios, K3 | Titan Krios, K3 | Titan Krios, K3 | Titan Krios, K3 | Titan Krios, K3 | Titan Krios, K3 | Titan Krios, K3 | Titan Krios, K3 | Titan Krios, K3 | Titan Krios, K3 | Titan Krios, K3 | Titan Krios, K3 | Titan Krios, K3 | Titan Krios, K3 |
| Magnification | 22,500× | 22,500× | 22,500× | 22,500× | 22,500× | 22,500× | 22,500× | 22,500× | 22,500× | 22,500× | 22,500× | 22,500× | 22,500× | 22,500× | 22,500× |
| Voltage (kV) | 300 | 300 | 300 | 300 | 300 | 300 | 300 | 300 | 300 | 300 | 300 | 300 | 300 | 300 | 300 |
| Dataset ID(s) | 20190816, 20191122 | 20190816, 20191122 | 20190816, 20191122 | 20190816, 20191122 | 20190816, 20191122 | 20190816, 20191122 | 20190816, 20191122 | 20190816, 20191122 | 20190816, 20191122 | 20190816, 20191122 | 20190816, 20191122 | 20190816, 20191122 | 20190816, 20191122 | 20200626 | 20200626 |
| Electron exposure (e–/Å²) | 33.960, 36.960 | 33.960, 36.960 | 33.960, 36.960 | 33.960, 36.960 | 33.960, 36.960 | 33.960, 36.960 | 33.960, 36.960 | 33.960, 36.960 | 33.960, 36.960 | 33.960, 36.960 | 33.960, 36.960 | 33.960, 36.960 | 33.960, 36.960 | 33.992 | 33.992 |
| Defocus range (μm) | -1.0 to -3.0 | -1.0 to -3.0 | -1.0 to -3.0 | -1.0 to -3.0 | -1.0 to -3.0 | -1.0 to -3.0 | -1.0 to -3.0 | -1.0 to -3.0 | -1.0 to -3.0 | -1.0 to -3.0 | -1.0 to -3.0 | -1.0 to -3.0 | -1.0 to -3.0 | -1.0 to -3.0 | -1.0 to -3.0 |
| Physical pixel size (Å) | 1.096 | 1.096 | 1.096 | 1.096 | 1.096 | 1.096 | 1.096 | 1.096 | 1.096 | 1.096 | 1.096 | 1.096 | 1.096 | 1.096 | 1.096 |
| Symmetry imposed | None (C1) | None (C1) | None (C1) | None (C1) | None (C1) | None (C1) | None (C1) | None (C1) | None (C1) | None (C1) | None (C1) | None (C1) | None (C1) | None (C1) | None (C1) |
| Initial particle images (no.) | 1889717 | 1889717 | 1889717 | 1889717 | 1889717 | 1889717 | 1889717 | 1889717 | 1889717 | 1889717 | 1889717 | 1889717 | 1889717 | 1473284 | 1473284 |
| Final particle images (no.) | 50776 | 114008 | 340359 | 21214 | 22586 | 104003 | 23775 | 32677 | 55876 | 325236 | 121796 | 128733 | 476440 | 341427 | 341427 |
| Map resolution (Å) FSC threshold = 0.143 | 4.03 | 3.65 | 3.29 | 3.67 | 3.52 | 3.56 | 4.50 | 4.23 | 3.83 | 4.14 | 3.86 | 3.88 | 3.15 | 3.09 | 3.58 |
| Map sharpening B factor (Å²) | 134.8 | 141.7 | 136.6 | 71.4 | 74.0 | 134.5 | 128.9 | 126.8 | 119.7 | 209.8 | 139.9 | 141.9 | 132.3 | 126.7 | 164.4 |
| **Model Refinement** | | | | | | | | | | | | | | | |
| Initial model used (PDB ID) | #3–4, #23 | #3–4, #23 | #3–4, #23 | #3–4, #23 | #3–4, #23 | #3–4, #23 | #3–4, #23 | #3–4, #23 | #3–4, #23 | 1JTH | #3–4 | #3–4 | #3–4 | #3–4, #28 | 1JTH |
| Model composition | | | | | | | | | | | | | | | |
| *Non-hydrogen atoms* | 42375 | 40071 | 25898 | 25042 | 25413 | 40590 | 42362 | 42422 | 25590 | 11330 | 29851 | 26763 | 29675 | 40475 | 13917 |
| *Protein residues* | 5376 | 4992 | 3205 | 3165 | 3206 | 5126 | 5354 | 5374 | 3204 | 1426 | 3713 | 3306 | 3505 | 4964 | 1760 |
| *Ligands* | 13 | 25 | 23 | 17 | 24 | 15 | 12 | 12 | 17 | 0 | 19 | 20 | 23 | 11 | 0 |
| B factors (Å²) | | | | | | | | | | | | | | | |
| *Protein* | 155.36 | 86.04 | 64.94 | 116.71 | 94.71 | 148.28 | 295.12 | 220.25 | 91.07 | 162.68 | 99.49 | 91.00 | 106.79 | 80.87 | 70.80 |
| R.M.S. deviations | | | | | | | | | | | | | | | |
| *Bond lengths (Å)* | 0.003 | 0.003 | 0.003 | 0.005 | 0.006 | 0.004 | 0.003 | 0.005 | 0.004 | 0.004 | 0.003 | 0.004 | 0.010 | 0.005 | 0.004 |
| *Bond angles (°)* | 0.575 | 0.619 | 0.649 | 1.065 | 1.147 | 0.990 | 0.702 | 1.089 | 0.799 | 0.678 | 0.666 | 1.062 | 1.681 | 1.043 | 0.556 |
| Validation | | | | | | | | | | | | | | | |
| *MolProbity score* | 1.77 | 1.57 | 1.38 | 1.68 | 1.57 | 1.67 | 1.92 | 1.88 | 1.75 | 2.05 | 1.52 | 1.69 | 1.42 | 1.74 | 1.56 |
| *Clashscore* | 8.61 | 5.20 | 2.90 | 5.75 | 4.10 | 6.98 | 14.16 | 9.43 | 7.33 | 9.14 | 4.80 | 6.39 | 3.62 | 4.94 | 8.66 |
| *Rotamer outliers (%)* | 0.04 | 0.47 | 0.29 | 0.56 | 1.10 | 0.30 | 0.50 | 1.12 | 0.73 | 2.44 | 0.34 | 0.28 | 0.94 | 2.09 | 0.54 |
| Ramachandran Statistics | | | | | | | | | | | | | | | |
| *Favored (%)* | 95.59 | 95.78 | 95.67 | 94.68 | 95.01 | 95.93 | 96.11 | 94.97 | 95.07 | 96.03 | 96.00 | 95.14 | 96.08 | 96.44 | 97.57 |
| *Allowed (%)* | 4.37 | 4.16 | 4.27 | 5.19 | 4.99 | 3.95 | 3.70 | 4.90 | 4.74 | 3.62 | 3.91 | 4.58 | 3.75 | 3.31 | 2.31 |
| *Outliers (%)* | 0.04 | 0.06 | 0.06 | 0.13 | 0.00 | 0.12 | 0.19 | 0.13 | 0.19 | 0.35 | 0.09 | 0.28 | 0.17 | 0.24 | 0.12 |

#### Supplementary Table 1 | Cryo-EM data, map, and model statistics

Data, map, and model statistics for the 25 NSF, sx20S, 21bin20S, 22bin20S, and α-SNAP—SNARE subcomplex models. See Supplementary Fig. 11 for corresponding FSC curves, Fourier sampling, and relative signal for all reconstructions listed here.

| Index | Model | Buffer Nucleotide | EDTA or Mg | Spire SNARE complex | SNARE in D1 pore | F engaged | A | B | C | D | E | F |
|---|---|---|---|---|---|---|---|---|---|---|---|---|
| 1 | 20s_sx1a-tet_atp-edta_c20-33_full_model.pdb | ATP | EDTA | syntaxin H3 SNARE tetramer | syntaxin-1a | ☑ | ADP | ATP | ATP | ATP | ATP | ATP |
| 3 | 20s_binary_atp-edta_c4-3-5-3_full_model.pdb | ATP | EDTA | ? | ? | ☑ | ADP | ATP | ATP | ATP | ATP | ATP |
| 4 | 20s_binary_atp-edta_c4-3-5-4_full_model.pdb | ATP | EDTA | ? | ? | ☐ | ADP | ATP | ATP | ATP | ATP | Apo |
| 5 | 20s_binary_atp-edta_c4-27-30-46_full_model.pdb | ATP | EDTA | Full 2:1 binary complex (21bin20S) | syntaxin-1a | ☑ | ADP | ATP | ATP | ATP | ATP | ATP |
| 6 | 20s_binary_atp-edta_c5-47_full_model.pdb | ATP | EDTA | Full 2:1 binary complex (21bin20S) | syntaxin-1a | ☐ | ADP | ATP | ATP | ATP | ATP | Apo |
| 7 | 20s_binary_atp-edta_c7-25-34-45_full_model.pdb | ATP | EDTA | Full 2:1 binary complex (21bin20S) | syntaxin-1a | ☑ | ADP | ATP | ATP | ATP | ATP | ATP |
| 8 | 20s_binary_atp-edta_c14-15-20-42_full_model.pdb | ATP | EDTA | Full 2:1 binary complex (21bin20S) | syntaxin-1a | ☑ | ADP | ATP | ATP | ATP | ATP | ATP |
| 9 | 20s_binary_atp-edta_c22-35-37-39_full_model.pdb | ATP | EDTA | Full 2:1 binary complex (21bin20S) | SNAP-25 | ☑ | ADP | ATP | ATP | ATP | ATP | ATP |
| 10 | 20s_binary_atp-edta_c23_full_model.pdb | ATP | EDTA | Full 2:1 binary complex (21bin20S) | syntaxin-1a | ☐ | ADP | ATP | ATP | ATP | ATP | Apo |
| 11 | 20s_binary_atp-edta_c31_full_model.pdb | ATP | EDTA | Full 2:1 binary complex (21bin20S) | SNAP-25 | ☐ | ADP | ATP | ATP | ATP | ATP | Apo |
| 12 | 20s_binary_atp-edta_c33_full_model.pdb | ATP | EDTA | Full 2:1 binary complex (21bin20S) | syntaxin-1a | ☑ | ADP | ATP | ATP | ATP | ATP | ATP |
| 14 | 20s_binary_atp-mg_c2-11_full_model.pdb | ATP | Mg | Full 2:2 binary complex (22bin20S) | syntaxin-1a | ☐ | ADP | ADP | ADP | ADP + Pi | ATP | ADP |
| 15 | 20s_binary_atp-mg_c3-5-6-9_full_model.pdb | ATP | Mg | Full 2:2 binary complex (22bin20S) | syntaxin-1a | ☐ | ADP | ADP + Mg | ADP + Pi + Mg | ADP + Pi + Mg | ATP + Mg | ADP |
| 16 | 20s_binary_atp-mg_c4-7-16-20-24_full_model.pdb | ATP | Mg | ? | ? | ☐ | ADP | ADP | ADP + Pi + Mg | ADP + Pi + Mg | ATP + Mg | ADP |
| 17 | 20s_binary_atp-mg_c4-7-16-20-24-m0-sO_full_model.pdb | ATP | Mg | ? | ? | ☑ | ADP | ADP + Pi | ADP + Pi | ADP + Pi + Mg | ATP + Mg | ATP |
| 18 | 20s_binary_atp-mg_c4-7-16-20-24-m0-sP_full_model.pdb | ATP | Mg | ? | ? | ☐ | ADP | ADP + Mg | ADP + Pi + Mg | ADP + Pi + Mg | ATP + Mg | ADP |
| 19 | 20s_binary_atp-mg_c8-14-21_full_model.pdb | ATP | Mg | Full 2:2 binary complex (22bin20S) | syntaxin-1a | ☐ | ADP | ADP + Pi + Mg | ADP + Pi + Mg | ADP + Pi + Mg | ATP + Mg | ADP |
| 20 | 20s_binary_atp-mg_c13_full_model.pdb | ATP | Mg | Full 2:2 binary complex (22bin20S) | syntaxin-1a | ☐ | ADP | ADP | ADP | ADP | ATP | ADP |
| 21 | 20s_binary_atp-mg_c15_full_model.pdb | ATP | Mg | ? | ? | ☐ | ADP | ADP | ADP | ADP | ATP | ADP |
| 22 | 20s_binary_atp-mg_c17_full_model.pdb | ATP | Mg | Full 2:2 binary complex (22bin20S) | ? | ☐ | ADP | ADP | ADP + Pi | ADP + Pi | ATP + Mg | ADP |
| 23 | 20s_binary_atp-mg_c19_full_model.pdb | ATP | Mg | ? | ? | ☐ | ADP | ADP + Pi | ADP + Pi | ADP + Mg | ATP | ADP |
| 24 | apo_binary_atp-mg_c3-24_full_model.pdb | ATP | Mg | None | None | N/A | ADP | ADP | ADP + Pi | ADP + Pi | ADP + Pi | ADP |
| 25 | apo_binary_atp-mg_c5-19_full_model.pdb | ATP | Mg | None | None | N/A | ADP | ADP | ADP + Pi | ADP + Pi | ADP | ADP |
| 26 | apo_binary_atp-mg_c6-11-13-17-18-22_full_model.pdb | ATP | Mg | None | None | N/A | ADP | ADP + Pi | ADP + Pi | ADP + Pi | ADP + Pi | ADP |
| 27 | 20s_binary-mini_atp-edta_c5-XXX-46_full_model.pdb | ATP | EDTA | Minimal 2:2 binary complex (min22bin20S) | SNAP-25 | ☐ | ADP | ATP | ATP | ATP | ATP | Apo |

**Supplementary Table 2 | Cryo-EM class reference**

Model details related to substrate engagement and nucleotide state for the 25 NSF, sx20S, 21bin20S, and 22bin20S models.

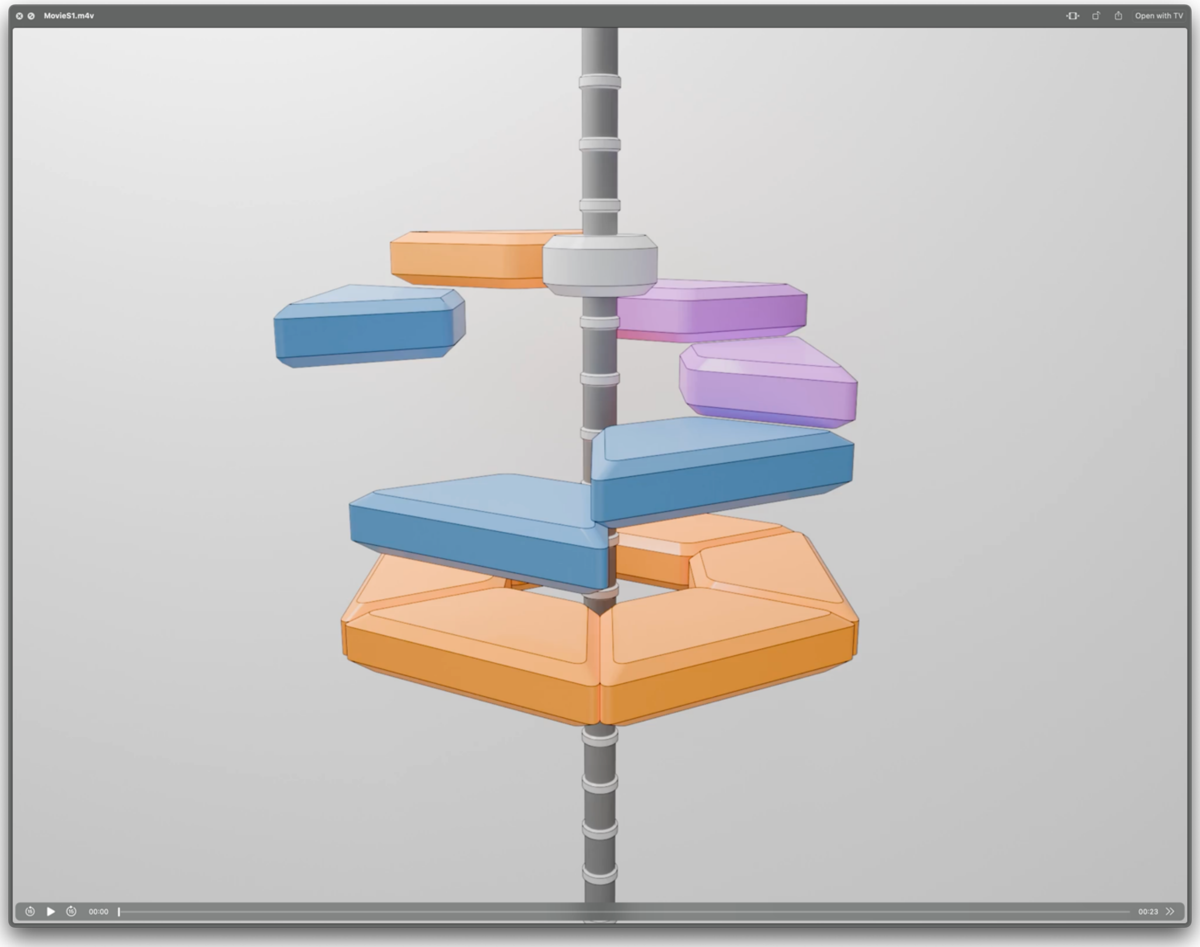

**Supplementary Movie 1 | A model of NSF motions and substrate translocation associated with nucleotide rebinding and hydrolysis.** A sequential hydrolysis model for SNARE complex disassembly by NSF inferred from comparison of different classes from non-hydrolyzing and hydrolyzing conditions. Following hydrolysis-independent substrate side loading, the D1 ring is found in a pre-hydrolysis state with ATP in protomer E of D1. Protomer F nucleotide exchange then enables ATP binding, re-engagement to both SNARE substrate, and the formation of the interprotomer interface with protomer E, triggering ATP hydrolysis in protomer E and substrate translocation. This is accompanied by $P_i$ and $Mg^{2+}$ release from down-ring protomers; in many structures of the different classes, this apparent release occurs between protomers C and B. The cycle then begins anew. Blue corresponds to ADP-bound, red to ATP-bound, purple to ADP•$P_i$ bound, and grey to nucleotide-free. See Fig. 10 for key frames and further detail.
